# HaploExplore, first software specifically designed for the detection of minor allele (MiA-) Haploblocks

**DOI:** 10.1101/2025.04.23.650206

**Authors:** Matilde Manetti, Samuel Hiet, Myriam Rahmouni, Jean-Louis Spadoni, Alice Dobiecki, Marco Lamanda, Maxime Tison, Taoufik Labib, Cristina Giuliani, Sigrid Le Clerc, Jean-François Zagury

**Author notes:** Corresponding author: Jean-François Zagury. The 2 first authors share an equal contribution for this work (Matilde Manetti, Samuel Hiet).

## Abstract

**Motivation:** Haplotype blocks in the genome are informative of evolutionary processes and they play a pivotal role in describing the genomic variability across human populations and susceptibility/resistance to diseases. Several software have been developed for haplotype blocks detection, but they do not distinguish between the impacts of major and minor SNP alleles. In this study, we present a powerful haploblock detection software, specifically designed for identifying haploblocks associated with SNP minor allele haploblocks (MiA-haploblocks). These haploblocks are particularly important as they can significantly influence phenotypic traits, offering a novel approach for studying genetic associations and complex traits.

**Results:** HaploExplore operates on VCF files containing phased data, exhibiting rapid processing times and generating user-friendly outputs. Its results are convergent for populations starting from 100 individuals. A comparative analysis of HaploExplore against other haploblock detection software revealed its superiority in terms of either simplicity, or flexibility, or speed, with the unique capability to target minor alleles. HaploExplore will be very useful for evolutionary genomics and for GWAS analysis in human diseases, given that the effects of genetic associations may accumulate within a specific haploblock.

## Introduction

Haplotypes are combinations of variant alleles with substantial linkage disequilibrium (LD) within a genomic region of a population [1]. The set of haplotypes found in a genomic region can delineate haplotype blocks, LD blocks or haploblocks, which provide valuable insights into evolutionary and recombinatory processes useful for applications ranging from population genetics to disease mapping [1,2]. LD block regions vary in size, with strong LD persisting within blocks until interruption, often due to recombination hotspots or population genetic phenomena [3,4]. Their patterns and position are shaped by recombination, mutation, selection, demography and various other evolutionary forces [5].

Haploblocks are indispensable in genomic research and genome-wide association studies (GWAS) due to their ability to represent high-LD regions, allowing the identification of genomic areas linked to complex traits and disorders [6]. Moreover, population differences in LD blocks may be one of the reasons for replication issues in GWAS across different populations. Analyzing haploblocks improves understanding of gene regulation, epistatic interactions, and evolutionary patterns [7]. By grouping associated variants, haploblocks simplify LD structures, and may help reduce noise in GWAS analyses, and improve the statistical power to identify causal variants [8]. They offer a framework for understanding how genetic variations collectively influence phenotypic outcomes and biological processes and provide insights into population-specific genetic selection, admixture events, and ancestral recombination patterns.

In GWAS, haploblocks may also capture the cumulative effects of variants in LD, offering a broader perspective than single genetic markers. For example, in a recent study, our group has identified a 1.9 megabase haploblock of 376 Single Nucleotides Polymorphisms (SNPs) associated with the allele HLA-B57:01 [9]. By exploring the haploblock structure “containing” this HLA allele, we were able to identify SNPs having cumulative effects on the phenotype within the haploblock, explaining the major signal observed for HLA-B*57:01 for HIV-1 elite control [10]. By focusing on haploblocks, researchers can thus uncover interdependencies and synergistic effects often missed in single-variant approaches, revealing critical factors in disease resistance and susceptibility [10].

A critical issue not addressed by existing software is the need to focus on minor alleles. One can reasonably conceive that some low frequency alleles have been selected in the course of evolution because they are associated to resistance to specific diseases, as has been described for the famous D32 mutant of the co-receptor CCR5 probably selected in Europeans for its better resistance to plague [11]. Conversely, some low frequency alleles may be selected during evolution due to their benefits but also increased susceptibility to certain diseases, as has been shown for the *β*^S^ allele (β-globin gene), that protects against malaria but acts as risk allele to sickle-cell disease [12]. Indeed, studies have shown that 1. minor alleles are more likely to be associated with disease susceptibility and resistance [13], and 2. disease-associated alleles are more likely to be low-frequency derived alleles than neutral expectations [14]. Focusing on minor alleles thus becomes a useful strategy to look for genetic markers of disease resistance/susceptibility. In order to address the need for a robust analysis of minor alleles in haploblocks, we have developed the HaploExplore software, specifically designed to facilitate such investigations. It works with phased genotypes and can handle very large genomic regions and datasets, involving thousands of SNPs, with the capacity to build haploblocks composed of the SNPs minor allele (MiA-haploblocks). It relies on explicit parameters such as Minor Allele Frequency (MAF), r^2^, D’, and carrier percentage (CP), which makes it suitable for a variety of applications. Its usefulness for genomic analyses is completed by simple output visualizations.

## 2. Material and Methods

### Test population

In this study, for sake of performance evaluation, we have used subgroups of a cohort genotyped with an Illumina chip, the DESIR (Data from an Epidemiological Study on Insulin Resistance Syndrome) cohort. It included 2,576 men and 2,636 women (aged 30–64 years at enrollment, 1994–1996) from the general French population [15] and a sub-cohort of approximately 1500 subjects was genotyped using the Illumina Infinium II HumanHap300 BeadChips (Illumina, San Diego, CA) [14]. For the present study, we randomly selected 500 individuals and focused on chromosome 22. Data was imputed and phased using the Michigan Imputation Server with the HRC r1.1 2016 (GRCh37/hg19) reference panel, imputation followed a quality filter of r^2^ > 0.3, and phasing was performed using Eagle v2.4 under the European (EUR) population model. After imputation, we obtained 125,956 SNPs on chromosome 22.

### Algorithm principle

The software processes a Variant Call Format (VCF) file [16] (compressed or not) and extracts the useful information (e.g. SNP-ID, position), and applies a filter based on the SNPs MAF. To speed up the analysis and reduce computational complexity, the algorithm splits the genome into smaller regions, with the region size being user-defined. To build these subregions the algorithm uses the sliding window approach, where overlapping subregions are created. The overlap size is equal to the maximum haploblock size parameter. Within each subregion, the SNPs are sorted in an ascending order of MAF, and the algorithm proceeds to scan the SNPs of the subregion according to this order. The algorithm starts from the first SNP of the ordered list - designated as the core SNP - serving as the anchor for the first haploblock. The coreSNP serves for identifying a group of SNPs whose minor alleles are genetically linked to its minor allele. The algorithm selects one by one the other SNPs (SNPtested) in the list and tries to add them in the haploblock according to the parameters. If the SNPtested is not too far from the coreSNP - based on the value of the maximum haploblock size parameter - then the algorithm checks if the MAF of the SNPtested is greater than the MAF cut threshold of the coreSNP. If so, the algorithm computes r^2^ and D’ between the coreSNP and the SNPtested, ensuring they meet the defined LD threshold. If the SNPtested reaches this step, the carrier percentage is computed in order to verify if the minor allele of the SNPtested is frequently co-occurred with the coreSNP’s minor allele, based on a predefined threshold. After the creation of haploblocks, the algorithm refines and corrects haploblocks by checking whether two consecutive SNPs within the block are too far apart, based on the position order. The default SNP gap threshold is 200 SNPs. Each genomic region is processed using one of three possible computational modes: the *Standard* mode, the *ListSNP* mode and the *Exhaustive* mode.

### Computational modes

The *Standard mode* constructs haploblocks iteratively. Each time a SNP does not fit in existing haploblocks, a new block is created using this SNP as the coreSNP. With the chosen LD parameters SNPs minor alleles are included in blocks if they meet the requested setting with regard to the coreSNP: LD between the SNP and the coreSNP, percentage carried by the coreSNP carriers, MAF percentage cut.

The *ListSNP* mode builds haploblocks based on a predefined list of SNPs, using the previously mentioned criteria. In this mode, haploblocks are initiated by SNPs present in the given list. These SNPs are considered as coreSNPs, and individual haploblocks are created for each SNP of the list. The user can set a first option in which SNPs from the list are used uniquely to generate the MiA-haploblocks, and no additional SNPs from the region will generate new haploblocks. Nevertheless, a second option allows the user to specify if additional SNPs in the region can also be used to create new haploblocks (thus resembling the standard mode). With a second parameter we can choose if the SNPs in the list can also participate to other haploblocks (option: Yes) or if the SNPs in the list cannot be added in others haploblocks and are thus only used as coreSNPs (option:No). Thus, the ListSNP mode is especially useful for focused analyses, such as studying specific loci of interest (e.g., HLA regions). It is possible to specify criteria such as LD thresholds, MAF, and CP cut-offs for refining haploblocks.

The *Exhaustive mode* aims to reconstruct haploblocks by iteratively scanning through all SNPs in a region. This iterative mode starts by assigning to each SNP its own haploblock. It employs LD and carrier percentage criteria to refine haploblocks while maintaining meaningful boundaries. This mode is particularly useful if the aim is to reconstruct haploblocks in regions with complex LD patterns or high genomic diversity.

### Main parameters

In the following paragraph coreSNP and SNPtested are referred as SNP1 and SNP2, respectively. Many parameters can be considered to identify haplotype blocks within a genomic region, among them, critical ones are : (1) MAF, SNPs are selected ensuring that the variants are common enough to be informative for block definition; (2) LD thresholds, such as r^2^ and D’ to make sure SNPs are sufficiently correlated with the coreSNP of the haploblock [17, 18]; (3) Carrier percentage (CP), this parameter is used to include a SNP2 minor allele in a haploblock when a sufficient proportion of individuals carrying a given SNP1 minor allele (coreSNP of the haploblock) also carry SNP2 minor allele. This ensures that SNPs are grouped with SNP1 when they are frequently observed together in individuals, allowing the haploblock to better represent their potential combined influence on genetic variation. The carrier percentage formula can be expressed as follows:

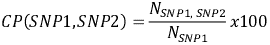

Where:

- *N*_SNPl, SNP2_ Is the number of individuals carrying the minor allele of SNP2 who also carry the minor allele of SNP1 (coreSNP);
- *N*_SNPl_ Is the total number of individuals carrying the minor allele of SNP1

Since we are referring to phased data, the percentage is computed separately for each chromatid:

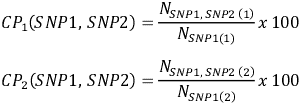

Where:

- *N*_SNPl, SNP2 (l)_ and *N*_SNPl, SNP2 (2)_are the number of individuals carrying the minor allele of SNP2 who also carry the minor allele of SNP1 on chromatid 1 and 2, respectively.
- *N*_SNPl(l)_and *N*_SNPl(2)_number of individuals carrying the minor allele of SNP1 on chromatid 1 and 2, respectively.

The overall exact carrier percentage across both chromatids is:

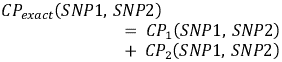

(4) The MAF cut threshold is defined as a relative threshold based on the minor allele frequency of the coreSNP to determine which SNPs can be included in a haploblock. To be clear:

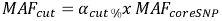

Where:

- *MAF*_*cut*_ is the minimum MAF required for a SNPtested to be included in the haploblocks
- α_*cut %*_is the MAF percentage cut, the actual tunable parameter in the software
- *MAF*_*coreSNP*_is the minor allele frequency of the coreSNP

Default values for these parameters are proposed and can be easily modified by the user (see Supplementary Materials).

#### 3. Results

### Impact of the population size

To evaluate the impact of sample size on haploblock detection and convergence, we used phased and imputed SNPs in the 35 Mb long chromosome 22 involving 125,956 SNPs data from the DESIR cohort of French individuals. Analyses were conducted on subsets of 25, 50, 100, 250 and 500 individuals, and two key outputs were assessed: the distribution of haploblock sizes in bp and the number of SNPs per haploblock (Figure 2A and 2B).

**Figure 1.**
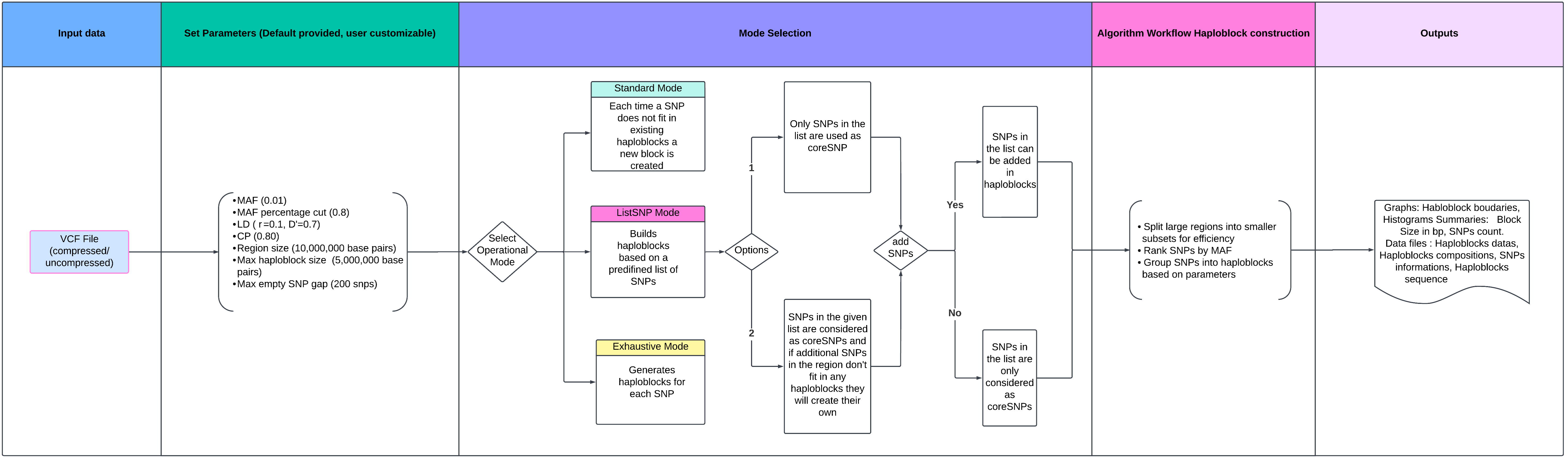
Workflow of HaploExplore, pipeline of the construction of the haploblocks

**Figure 2.**
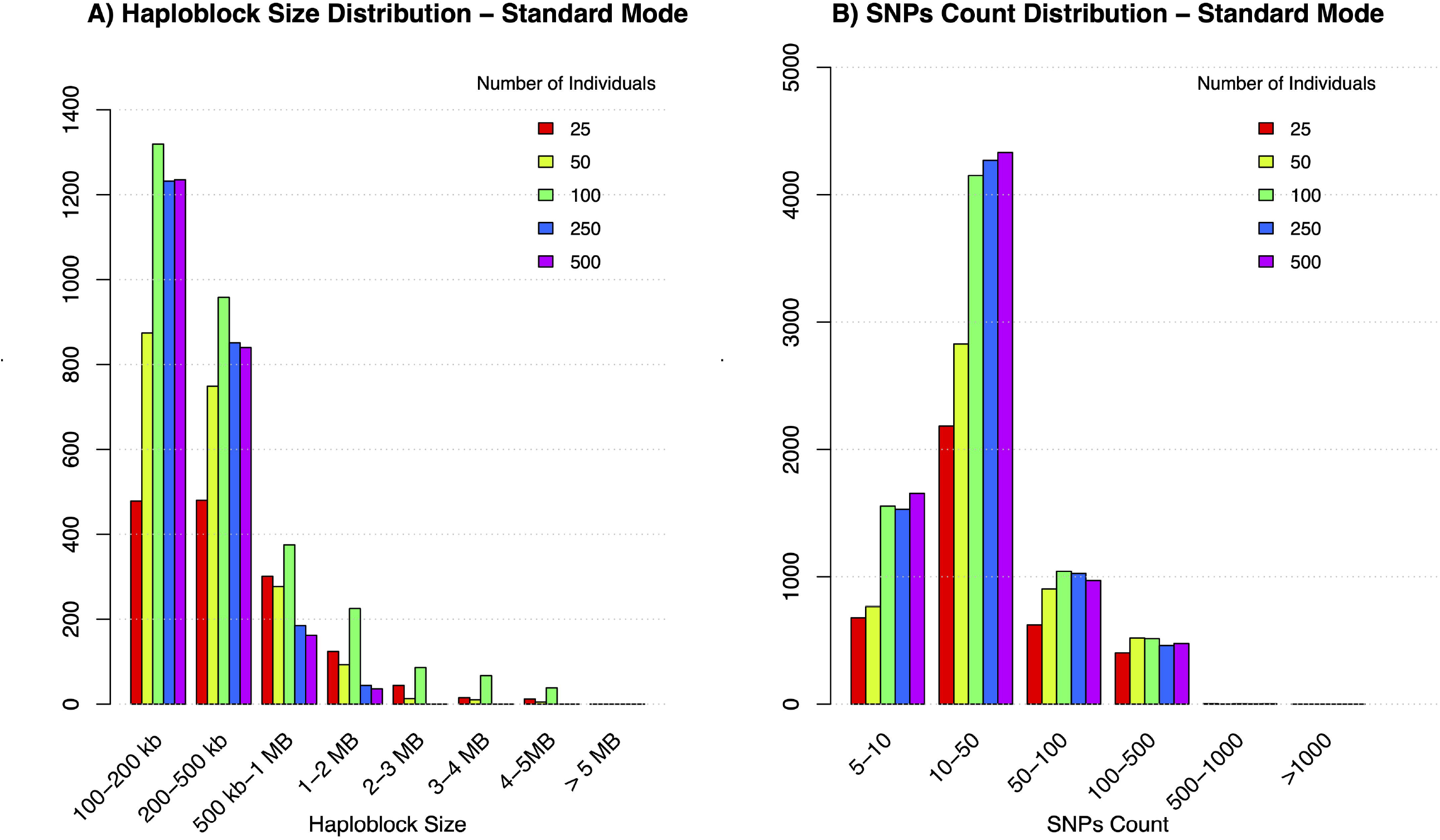
Haploblock and SNP Count Distributions for Standard Mode. (A)□Distribution of haploblock sizes across various sample sizes from the DESIR cohort. The x-axis displays haploblock size categories, ranging from 100 kb to over 5 MB, while the y-axis represents the number of haploblocks observed in each size category. Bar plots are used to show the distribution for each sample size (25, 50, 100, 250, 500 individuals), with distinct colors representing different sample sizes. The reduced plot excludes the smallest haploblock size category (0–100 kb) to enhance the visualization of larger haploblocks. (B)□Distribution of SNP counts per haploblock across various sample sizes in the analysis. The x-axis shows SNP count categories (ranging from 5 to >1,000 SNPs per haploblock), while the y-axis represents the number of haploblocks observed. Bar plots are used to display the results for all sample sizes (25, 50, 100, 250, 500 individuals), with each sample size represented by a unique color. The reduced plot excludes the categories 1, 2, 3, and 4 SNPs to provide a clearer view of the distribution for haploblocks with higher SNP counts. Both panels are based on data from□chr 22□(125,956 SNPs in 35 MB) using the□standard mode□of HaploExplore. Parameters include: LD thresholds (r^2^ ≥ 0.1, D′ ≥ 0.7),□MAF ≥ 0.8,□%carrier threshold ≥ 80%, region size of 10,000,000 base pairs□with a region overlap of□5,000,000 base pairs, and a maximum empty gap of□200 SNPs.

Results demonstrate that smaller sample sizes (between 25 and 100 individuals) produce more variable outputs, with wider fluctuations in both haploblock size and SNP counts (Figure 2A and 2B). This variability reflects a lack of convergence, likely driven by insufficient sample representation of linkage disequilibrium patterns. The results indicate a clear trend toward convergence, with diminishing changes beyond 100–250 individuals (*Figure 2A; 2B*). Additional details, including extended tables and figures supporting these findings, are provided in the Supplementary Materials. A simplified visualization of the results is presented in *Figure 2* by excluding the 0–100 kb haploblock size category and the categories 1, 2, 3, and 4 SNPs per haploblock. This adjustment improves clarity and focuses on the most informative haploblocks, particularly those of larger sizes and SNP counts. This approach has been applied for all the plots of the text to ease the visualization of results and emphasize the key trends and patterns relevant to the performed analyses. For completeness, the full results —including all haploblock size and SNP count categories— are provided in the Supplementary Materials. For these analyses, we have employed the *Standard Mode* for haploblock detection, with the default’s parameters. While the computational speed analysis will be addressed later, these first findings highlight the importance of an adequate sample size to achieve convergence and reliable detection of haploblock boundaries.

### Speed

The computational efficiency of a software is always an important factor, given the challenges posed by large sample sizes and extensive datasets. We evaluated the running time of the three processing modes—ListSNP, Standard, and Exhaustive—across varying sample sizes (25, 50, 100, 250 and 500 individuals) (Figure 3; Table S4). The running times were quite reasonable since they were in terms of minutes to hours according to the mode. For instance, for both phased and imputed data, the *ListSNP* mode with a list of 500 SNPs exhibited running times of a few minutes across all sample sizes (chromosome 22: 125,956 SNPs in 35 Mb) with only 2 minutes for 25 individuals and 6 minutes for 500 individuals (Table S4). By contrast, the exhaustive mode, which computes haploblocks for all SNPs in a region without relying on predefined lists, was significantly slower due to the increased computational demand, always considering the same chromosome 22. This mode processes a much larger number of SNPs, consequently, the exhaustive mode had running times ranging from 263.41 minutes for 25 individuals to 1215 minutes (around 20 hours) for 500 individuals over the chromosome 22.

**Figure 3.**
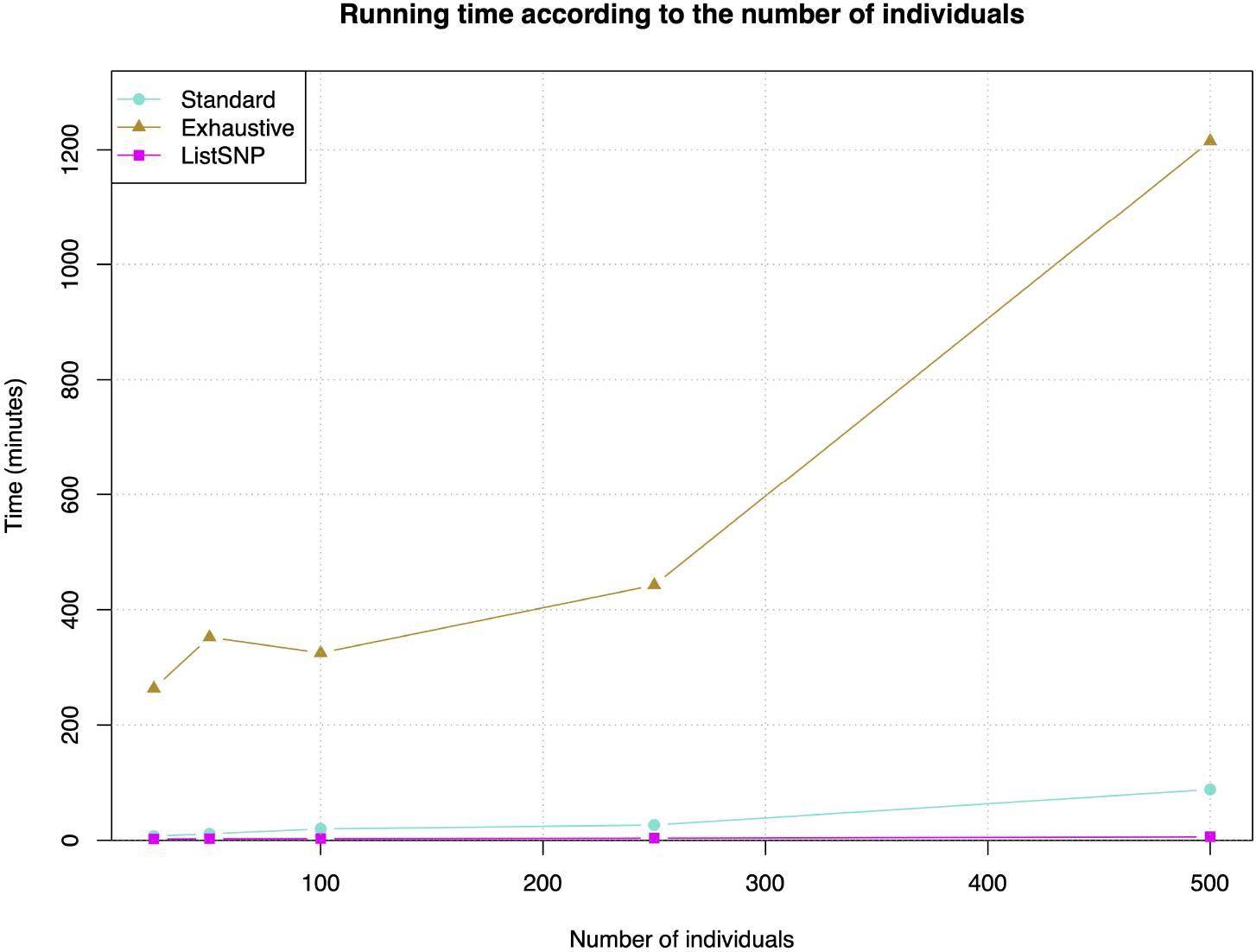
Running time of HaploExplore across different population sizes. (25, 50, 100, 250, and 500 individuals) for the Standard mode with LD, Exhaustive mode, and ListSNP mode on chr 22 - size of 35 Mb - 125,956 SNPs.

The S*tandard mode* was slower than the *ListSNP* mode but considerably faster than the Exhaustive mode. These results illustrate the trade-offs between computational speed and analysis scope. *ListSNP* is ideal for analyses targeting a specific set of SNPs, offering high efficiency and speed. However, the *Exhaustive* mode remains indispensable for comprehensive studies that require analyzing all SNPs in a genomic region. The choice of the mode depends on the scope of the study of interest and available computational resources.

### 3.1. Comparison with other haploblock detection software

Several computational tools exist for haploblock detection, each employing distinct methodologies and having differences in parameter flexibility. In Table 1, we compared their algorithm, parameters, and other features with those of HaploExplore.

**Table 1.**
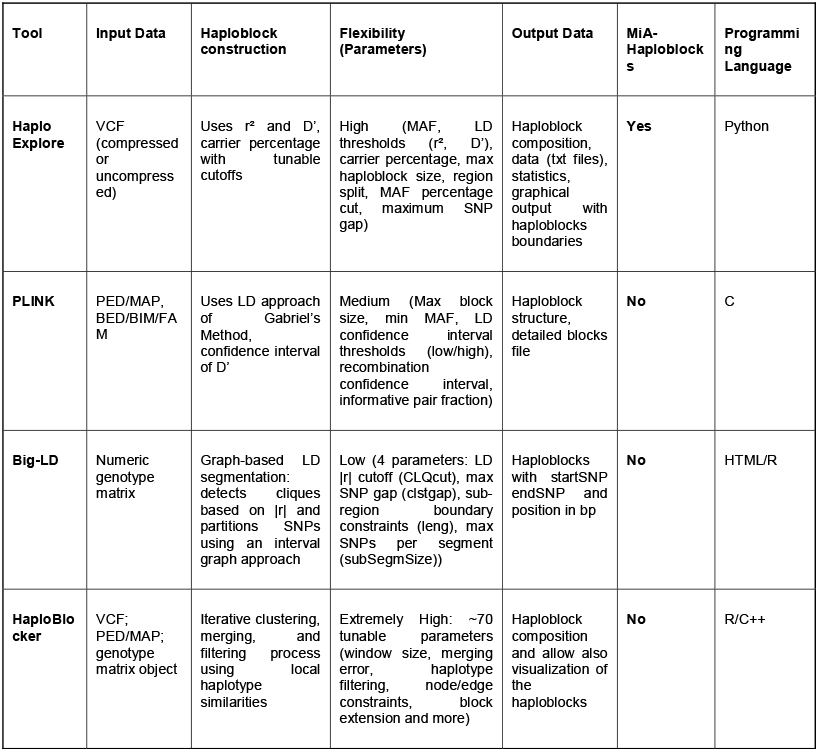
Comparison of Haploblock Detection Tools: Summary of key features across different haploblock detection tools, including input data formats, haploblock construction methods, parameter flexibility, output types, and support for MiA-haploblocks and programming language.

## Discussion

A new tool for haploblock detection has been developed, specifically designed to target minor alleles. It provides flexibility and precision by implementing the definition of haploblocks based on multiple biological and statistical parameters, including MAF, LD measures (r^2^ and D’), and carrier percentages. Its direct processing of standard VCF files ensures compatibility with widely used genomic data. HaploExplore manages effectively large genomic datasets by partitioning data into smaller genomic regions thereby allowing computational efficiency. It has the capacity to analyze an entire chromosome (more than 125,000 SNPs) in less than 10 mn.

Its modes of operation - *Standard Mode, ListSNP Mode and Exhaustive Mode* - provide diverse approaches to build and refine haploblocks based on user-defined thresholds and specific research objectives. The *Exhaustive* and *ListSNP* modes are particularly adapted for exploratory analyses and targeted research on small genomic regions. The *Exhaustive* mode is especially beneficial when researchers aim to analyze each single nucleotide polymorphism individually within a biologically relevant region, while the *ListSNP* mode is tailored for scenarios where a predefined list of SNPs is prioritized for MiA-haploblock analyses of specific genomic loci. In contrast, the *Standard* mode is designed for the analysis of entire genomes and the identification of haploblocks across extensive regions with varying SNP densities, making it the preferred choice for comprehensive genome exploration due to its scalability with increasing sample sizes.

The findings from our analysis of HaploExplore highlight the critical impact of sample size on haploblock detection and the robustness of different detection modes. Our results indicate that increasing the population size to 100 individuals or higher significantly enhances convergence, leading to more stable and reproducible haploblock distributions. The computational performance analysis highlights the scalability of HaploExplore. While the *Exhaustive* mode remains the most time and memory consuming, the *Standard* and *ListSNP* modes offer more efficient alternatives, with execution times remaining reasonable even for large genomic datasets. Different haploblock detection tools employ distinct methodologies, affecting their ability to capture MiA-haploblocks. PLINK and Big-LD define haploblocks based on traditional LD measures (such as D’, r^2^), making them fitted for common haplotype structures but limiting their ability to detect MiA-haploblocks. HaploBlocker, using sequence haplotype similarity-based clustering, prioritizes long, stable haplotypes but does not focus on minor allele inheritance patterns. HaploExplore uniquely integrates LD thresholds with a carrier percentage criterion, allowing the identification of MiA-haploblocks, which are crucial for studying genetic associations and complex traits. These haploblocks capture co-inherited minor alleles that may influence disease susceptibility and trait variability - patterns often missed by LD -based methods. Moreover, the software provides an interactive environment, allowing users to visualize and adjust parameters before each analysis, improving usability and supporting exploratory analysis.

While HaploExplore has demonstrated robust performance and speed in haploblock detection in its current form, some improvements could increase its efficiency and applicability. Notably we plan to optimize the software to improve speed and memory management by implementing its core algorithms in C++ (right now it is in Python). This transition will certainly reduce computational time significantly, making large-scale genomic studies even more accessible. To conclude, HaploExplore is a sophisticated and flexible software solution for haploblock detection and analysis, with a focus on minor alleles, offering a wide range of strengths that make it well-suited for genomic research.

### Web resources - Software Availability

We have developed an interactive web application using Streamlit, allowing users to run the software with a user-friendly interface. Default parameters are pre-set but can be adjusted directly from the application. A detailed description of the parameters, input, and output files is provided in the ReadMe.txt file.

## Data Availability Statement

The genomic data used in this article have been provided by the ICGH Consortium [10]. As genomic data, they cannot be transferred to the public but can be obtained under specific agreements upon contacting the coordinator of the ICGH Consortium (Dr Paul Mc Laren).

## Acknowledgments

The Laboratory GBCM and the Foundation Jean Dausset-CEPH are grateful to the program Mécénat-Santé of Mutuelles AXA for funding this research.

## Reference

1. Gabriel, Stacey B., Stephen F. Schaffner, Huy Nguyen, Jamie M. Moore, Jessica Roy, Brendan Blumenstiel, and others, ‘The Structure of Haplotype Blocks in the Human Genome’, Science, 296/5576 (2002), 2225–29

2. Sabeti, Pardis C., David E. Reich, John M. Higgins, Haninah Z. P. Levine, Daniel J. Richter, Stephen F. Schaffner, and others, ‘Detecting Recent Positive Selection in the Human Genome from Haplotype Structure’, Nature, 419/6909 (2002), 832–37

3. Jeffreys, Alec J., Liisa Kauppi, and Rita Neumann, ‘Intensely Punctate Meiotic Recombination in the Class II Region of the Major Histocompatibility Complex’, Nature Genetics, 29/2 (2001), 217–22

4. Stephens, Matthew, and Paul Scheet, ‘Accounting for Decay of Linkage Disequilibrium in Haplotype Inference and Missing-Data Imputation’, The American Journal of Human Genetics, 76/3 (2005), 449–62

5. McVean, Gilean A. T., Simon R. Myers, Sarah Hunt, Panos Deloukas, David R. Bentley, and Peter Donnelly, ‘The Fine-Scale Structure of Recombination Rate Variation in the Human Genome’, Science, 304/5670 (2004), 581–84

6. Cuyabano, Beatriz Cd, Guosheng Su, and Mogens S. Lund, ‘Selection of Haplotype Variables from a High-Density Marker Map for Genomic Prediction’, Genetics Selection Evolution, 47/1 (2015), 61

7. Traherne, J. A., ‘Human MHC Architecture and Evolution: Implications for Disease Association Studies’, International Journal of Immunogenetics, 35/3 (2008), 179–92

8. Karkar, Slim, Claire Dandine-Roulland, Jean-François Mangin, Yann Le Guen, Cathy Philippe, Jean-François Deleuze, and others, ‘Genome-Wide Haplotype Association Study in Imaging Genetics Using Whole-Brain Sulcal Openings of 16,304 UK Biobank Subjects’, European Journal of Human Genetics, 29/9 (2021), 1424–37

9. ahmouni, Myriam, Lorenzo De Marco, Jean-Louis Spadoni, Maxime Tison, Raissa Medina-Santos, Taoufik Labib, and others, ‘The HLA-B*57:01 Allele Corresponds to a Very Large MHC Haploblock Likely Explaining Its Massive Effect for HIV-1 Elite Control’, Frontiers in Immunology, 14 (2023), 1305856

10. Rahmouni, Myriam, Sigrid Le Clerc, Jean-Louis Spadoni, Taoufik Labib, Maxime Tison, Raissa Medina-Santos, and others, ‘Deep Analysis of the Major Histocompatibility Complex Genetic Associations Using Covariate Analysis and Haploblocks Unravels New Mechanisms for the Molecular Etiology of Elite Control in AIDS’, BMC Immunology, 26/1 (2025), 1

11. Duncan, S R, ‘Reappraisal of the Historical Selective Pressures for the CCR5-32 Mutation’, Journal of Medical Genetics, 42/3 (2005), 205–8

12. Laval, Guillaume, Stéphane Peyrégne, Nora Zidane, Christine Harmant, François Renaud, Etienne Patin, and others, ‘Recent Adaptive Acquisition by African Rainforest Hunter-Gatherers of the Late Pleistocene Sickle-Cell Mutation Suggests Past Differences in Malaria Exposure’, The American Journal of Human Genetics, 104/3 (2019), 553–61

13. Kido, Takashi, Weronika Sikora-Wohlfeld, Minae Kawashima, Shinichi Kikuchi, Naoyuki Kamatani, Anil Patwardhan, and others, ‘Are Minor Alleles More Likely to Be Risk Alleles?’, BMC Medical Genomics, 11/1 (2018), 3

14. Lachance, Joseph, ‘Disease-Associated Alleles in Genome-Wide Association Studies Are Enriched for Derived Low Frequency Alleles Relative to HapMap and Neutral Expectations’, BMC Medical Genomics, 3/1 (2010), 57

15. Limou, Sophie, Cédric Coulonges, Joshua T. Herbeck, Daniëlle Van Manen, Ping An, Sigrid Le Clerc, and others, ‘Multiple-Cohort Genetic Association Study Reveals CXCR6 as a New Chemokine Receptor Involved in Long-Term Nonprogression to AIDS’, The Journal of Infectious Diseases, 202/6 (2010), 908–15

16. Danecek, Petr, Adam Auton, Goncalo Abecasis, Cornelis A. Albers, Eric Banks, Mark A. DePristo, and others, ‘The Variant Call Format and VCFtools’, Bioinformatics, 27/15 (2011), 2156–58

17. Hill, W. G., and Alan Robertson, ‘Linkage Disequilibrium in Finite Populations’, Theoretical and Applied Genetics, 38/6 (1968), 226–31

18. Lewontin, R. C., ‘The Interaction of Selection and Linkage. I. General Considerations; Heterotic Models’, Genetics, 49/1 (1964), 49–67

