## Supplementary Materials for "HaploExplore, first software specifically designed for the detection of minor allele (MiA-) Haploblocks"

1. **Method**

*Default settings*

The software supports adjustable parameters, including: LD thresholds (default r² =0.1, D′=0.7), carrier percentage cut-off (default 0.8), MAF percentage cut (default 0.8), maximum SNP gap within a block (default 200 SNPs), maximum haploblock size (default 5M base pairs), region size for splitting datasets (default 10M base pairs), while the minimum MAF threshold to consider a SNP is 1% (default value).

Carrier Percentage plays a crucial role in detecting haploblocks of minor alleles even when traditional linkage disequilibrium measures, such as r², are low (e.g., 0.1) but D′ remains high (e.g., 0.7). Unlike LD-based approaches that rely on strong statistical correlations, CP directly assesses the co-occurrence of minor alleles within individuals, ensuring that SNP2 is included in a haploblock with SNP1 when they frequently appear together. By focusing on the actual presence of alleles in individuals rather than just correlation values, CP allows the detection of weaker genetic associations that might otherwise be missed due to a low r², this is particularly useful when SNPs are in high D' but low r². The default value for the carrier percentage threshold has been set at 80%. This choice is confirmed by both biological and statistical considerations. The haploblock should include SNPs whose minor alleles sufficiently cover the coreSNP minor allele to help explain the genetic association detected in statistical analyses. An 80% carrier percentage ensures that SNPs in the haploblock are strongly linked to the coreSNP. If a causal SNP is located within the haploblock of a coreSNP identified through statistical screening, we expect that the majority of individuals carrying the coreSNP minor allele also carry the minor allele of the causal SNP.

Setting the haploblock MAF percentage cut ($\alpha_{cut \%}$) at 0.8 of the coreSNP’s MAF ensures that only SNPs with a MAF of at least 80% of the coreSNP’s MAF are considered for inclusion. This allows SNPs with slightly lower frequencies than the coreSNP to still be part of the haploblock and contribute to explaining the statistical impact associated with the coreSNP. For example, if the coreSNP has a MAF of 0.4, only SNPs with a MAF of at least 0.32 will be evaluated for inclusion.

From our preliminary tests, the maximum SNP gap value of 200 seems to be a reasonable value which allows us to generate more compact haploblocks.

For efficiency reasons, the genome is cut in successive parts to explore the haploblocks, and we observed that the optimal region size to cut the genome was twice the maximal haploblock size.

*Output Data*

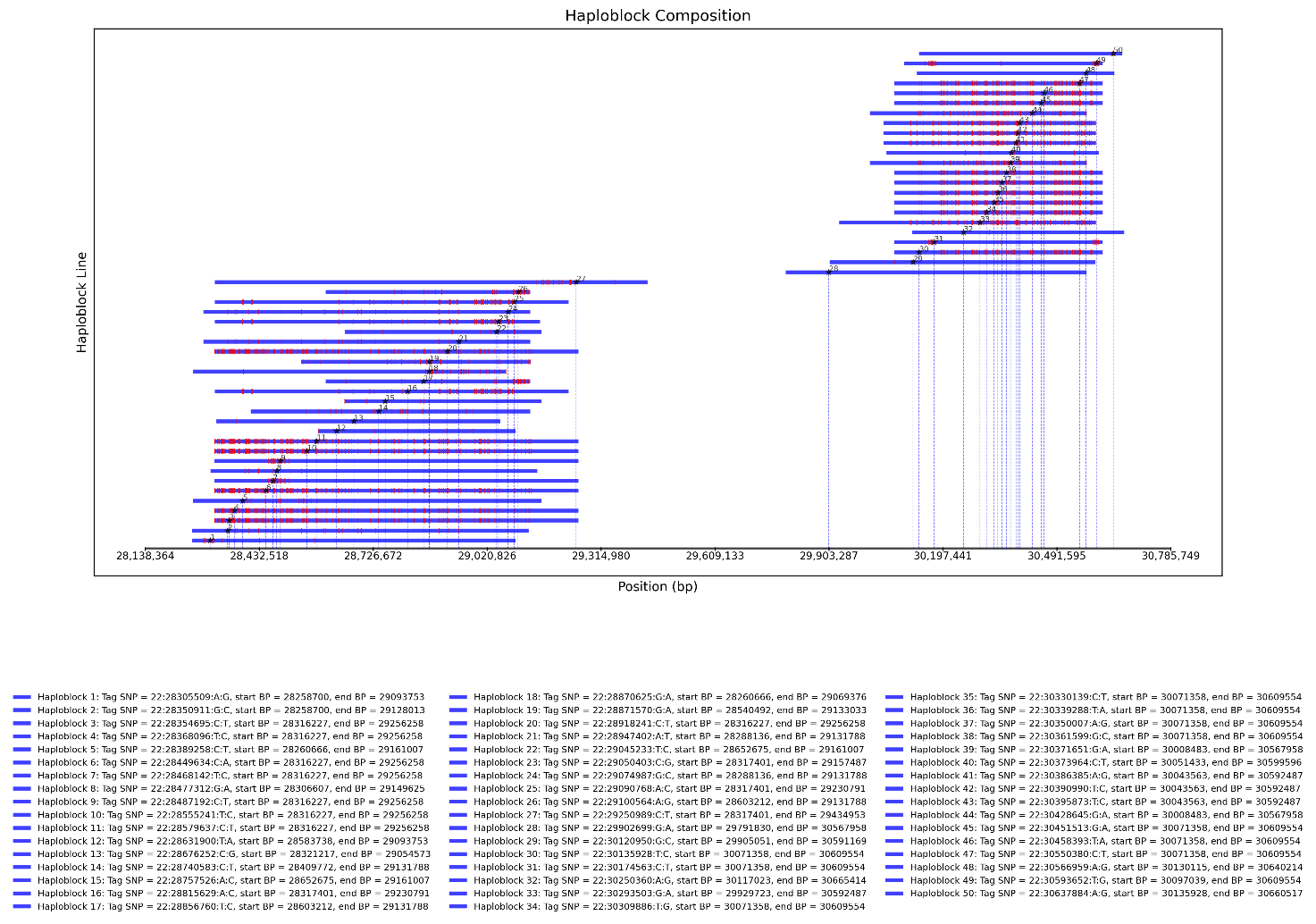

**Figure S1 Graphical Representations of Haploblock Boundaries Using List SNPs Mode.** *Haploblock boundaries are identified using the List SNPs mode. There were 1000 SNPs in the predefined SNP list, and they serve as coreSNP to build new haploblocks and are marked by a star. SNPs in red indicate those with a Minor Allele Frequency within ±10% of the MAF of the corresponding coreSNP (to visualize some kind of haplotype). The remaining SNPs within the haploblock are depicted in blue. This representation highlights the relationship between coreSNPs, surrounding SNPs, and the overall haploblock structure. The x-axis represents the SNP positions in base pairs (bp), while the y-axis corresponds to haploblock lines. The software creates more pdf files, for each pdf it creates a graph with maximum 50 haploblocks, the haploblocks are plotted according to the order of the list when they are generated, this is the first plot that we obtain from the analysis.*

**2. Results**

***Impact of population size***

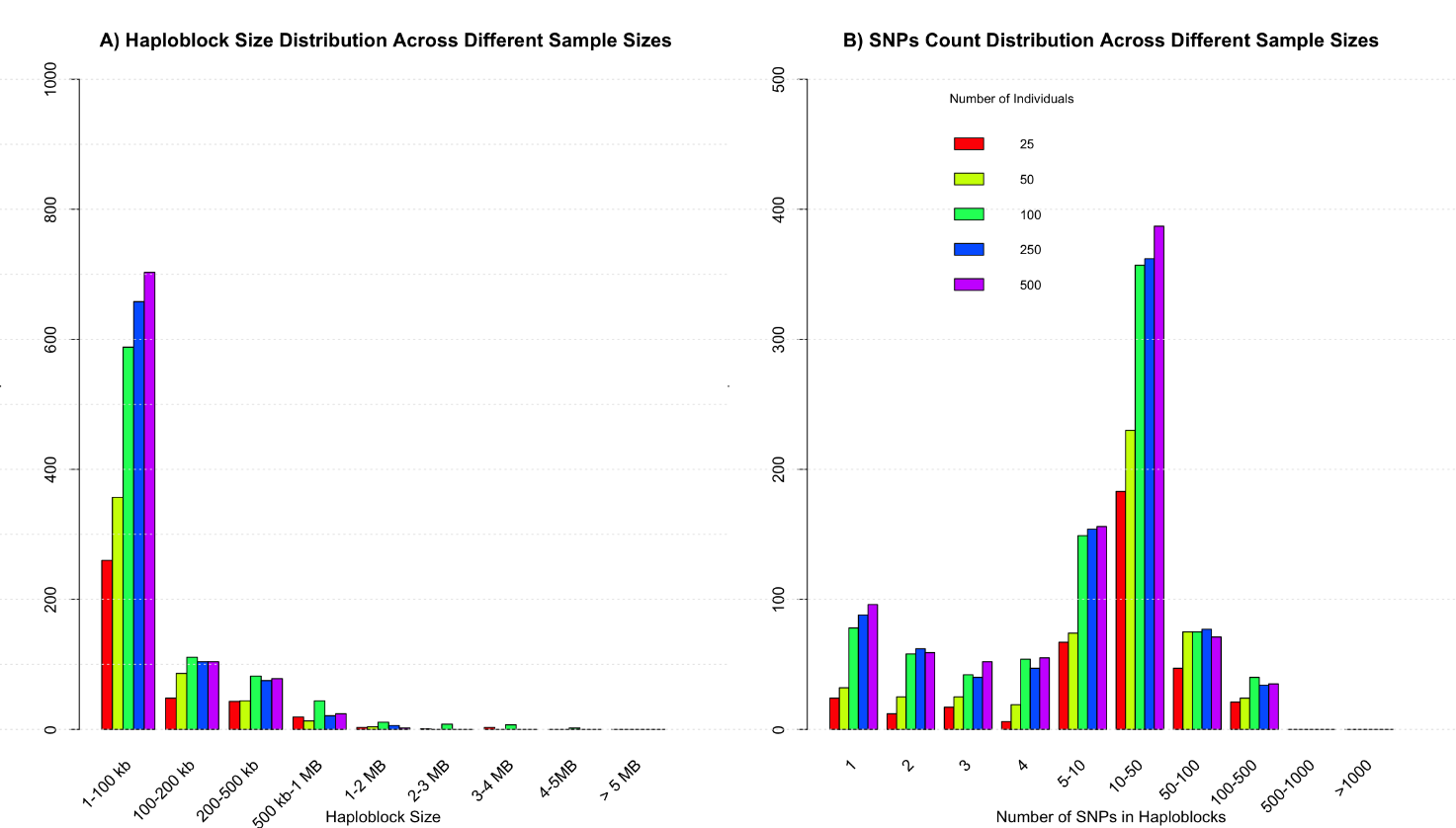

**Figure S2** **(A)** Distribution of haploblock sizes across various sample sizes from the DESIR cohort. The x-axis displays haploblock size categories, ranging from 1–100 kb to over 5 MB, while the y-axis represents the number of haploblocks observed in each size category. Bar plots are used to show the distribution for each sample size (25, 50, 100, 250, 500 individuals), with distinct colors representing different sample sizes. The results highlight how larger sample sizes (250–500 individuals) lead to a more stable and reliable representation of haploblock size distributions. **(B)** Distribution of SNP counts per haploblock across various sample sizes in the analysis. The x-axis shows SNP count categories (ranging from 1 to >1000 SNPs per haploblock), while the y-axis represents the number of haploblocks observed. Bar plots are used to display the results for all sample sizes (25, 50, 100, 250, 500 individuals), with each sample size represented by a unique color. Both panels are based on data from chr 22 using the standard mode of HaploExplore. Parameters include: LD thresholds (r² ≥ 0.1, D′ ≥ 0.7), MAF ≥ 0.8, %carrier threshold ≥ 80, region size of 10,000,000 base pairs with a region overlap of 5,000,000 base pairs, and a maximum empty gap of 200 SNPs.

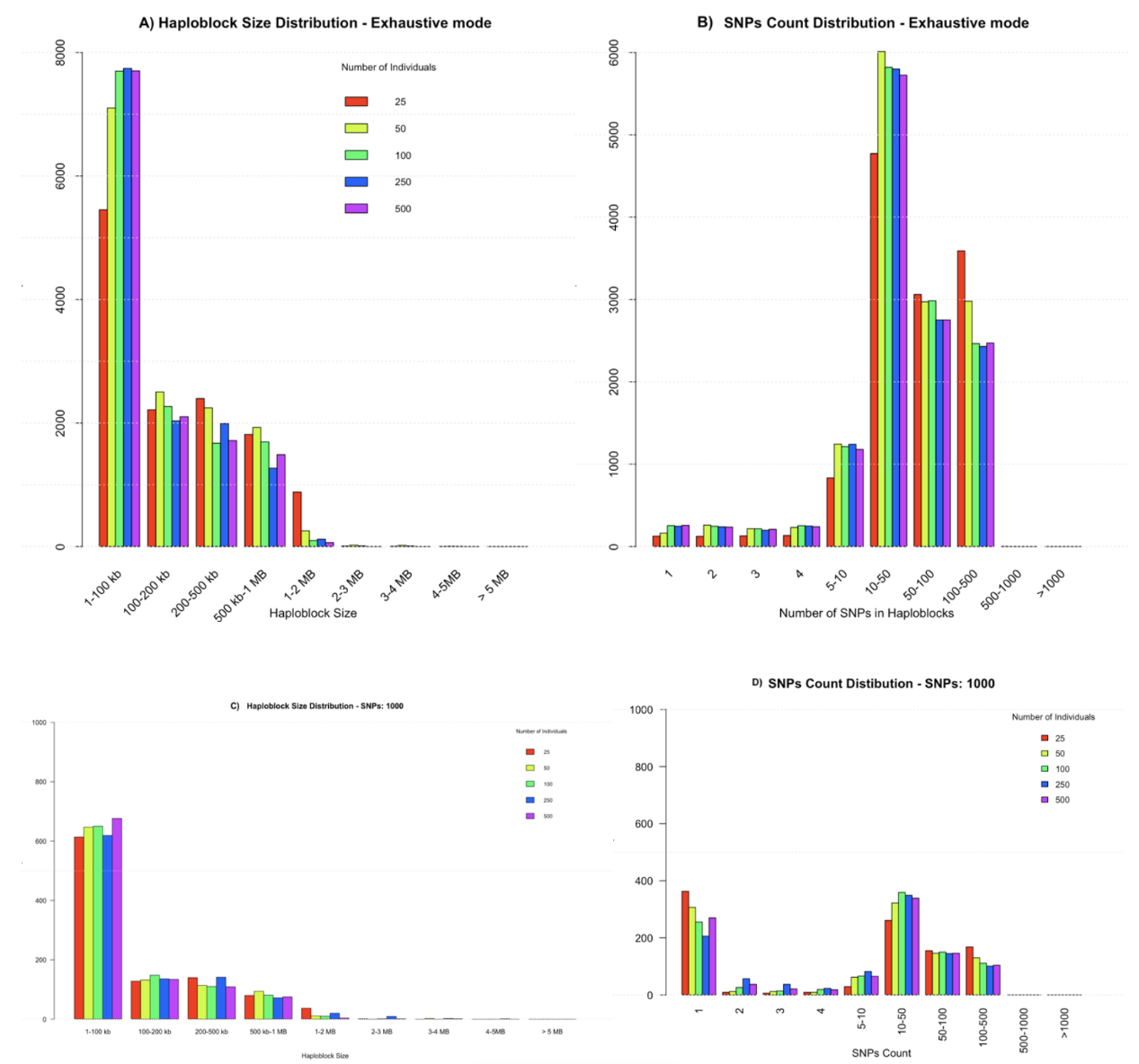

**Figure S3 Haploblock and SNP Count Distributions for Different Modes.** **(A)** Haploblock size distribution using the Exhaustive mode. **(B)** SNP count distribution using the Exhaustive mode. **(C)** Haploblock size distribution using the List SNPs mode. **(D)** SNP count distribution using the List SNPs mode, with the number of SNPs in the predefined list set to 1,000. These bar plots illustrate how haploblock sizes and the number of SNPs per haploblock vary with increasing sample sizes (25, 50, 100, 250, 500 genotyped individuals of the DESIR cohort). Each plot uses a smaller genomic region of 5 Mb of chromosome 22 for the analysis, with parameters: LD thresholds (r² ≥ 0.1, D′ ≥ 0.7), MAF ≥ 0.8, %carrier threshold ≥ 80, region size of 5,000,000 base pairs, and a maximum empty gap of 200 SNPs. These visualizations highlight the differences between modes and how sample size impacts haploblock structure and SNP distribution.

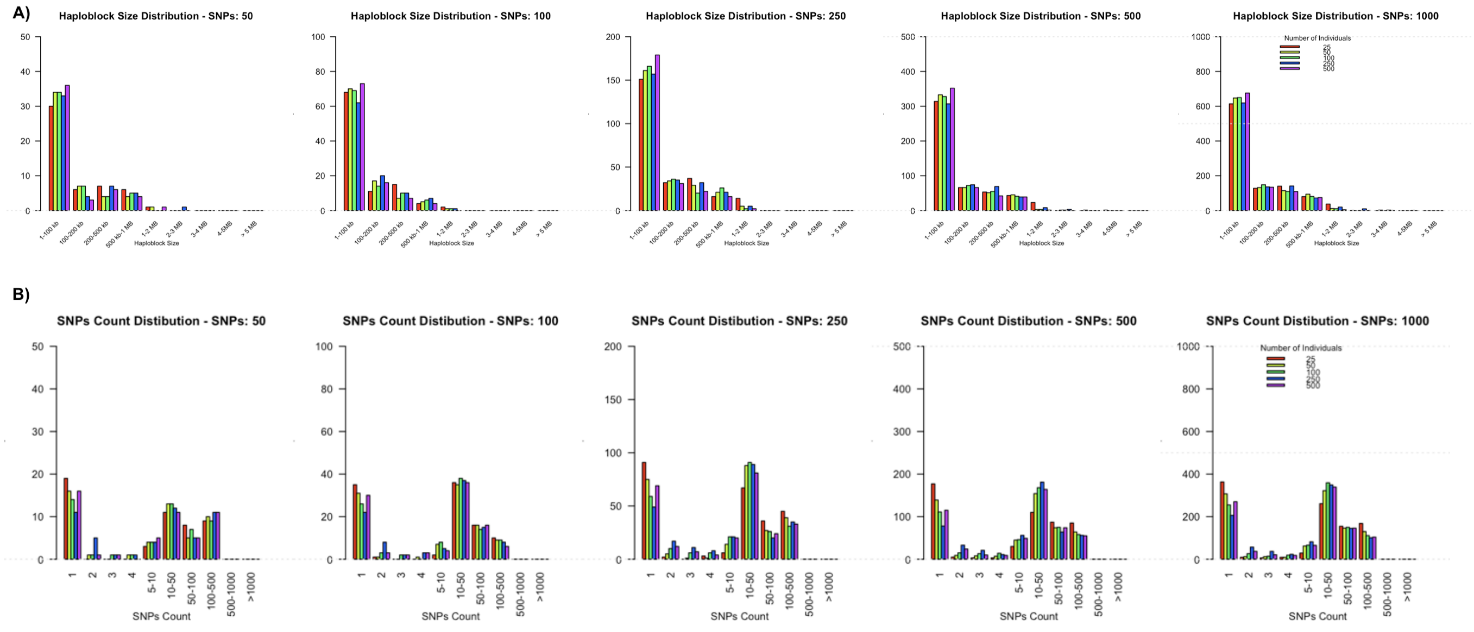

**Figure S4** **Haploblock Size and SNP Count Distributions for List SNPs Mode Across Different Predefined SNP Lists.** *Different bar plots for haploblock size and SNP count distributions generated using the List SNPs mode, with predefined SNP lists containing 50, 100, 250, 500, and 1,000 SNPs, with the option for which we have that the number of the SNPs in the list corresponds to the number of the haploblocks detected. The SNPs of the lists were chosen randomly from a subregion of the chromosome 22 (between positions 16,554,711 bp – 21,512,333 bp) and the input data is from the whole chromosome 22 in genotyped individuals from the DESIR cohort* **(A)** *Haploblock Size Distribution, five separate plots show the distribution of haploblock sizes for each predefined SNP list size. The x-axis displays haploblock size categories (ranging from 1–100 kb to over 5 MB), while the y-axis represents the number of haploblocks observed for each category. Distinct colors are used to represent different sample sizes (25, 50, 100, 250, 500 individuals).* **(B)** *SNP Count Distribution, five plots show the distribution of SNP counts per haploblock for each predefined SNP list size. The x-axis shows SNP count categories (ranging from 1 to >1,000 SNPs per haploblock), and the y-axis represents the number of haploblocks observed. Each sample size is represented by a unique color in the bar plots. These visualizations are based on a smaller genomic region of 5 Mb and were generated using the following parameters: LD thresholds (r² ≥ 0.1, D′ ≥ 0.7), MAF ≥ 0.8, %carrier threshold ≥ 80, region size of 5,000,000 base pairs, and a maximum empty gap of 200 SNPs.*

***Convergency***

To assess the stability of haploblock detection across different sample sizes, we analyzed the number of haploblocks identified and the corresponding genomic coverage as the sample size increased for the Standard mode with default parameters. The results indicate a clear trend toward convergence, with diminishing changes beyond 100–250 individuals.

**Table S1** Impact of sample size on haploblocks detection: number of haploblocks identified across different sample sizes of genotyped individuals of the DESIR cohort, with relative change indicating the increase in haploblock count as sample size grows

| **Sample Size** | **Haploblocks** | **Relative Change** |
| --- | --- | --- |
| 25 | 4,660 | - |
| 50 | 5,985 | +22.1% |
| 100 | 9,655 | +38.0% |
| 250 | 10,031 | +3.7% |
| 500 | 10,295 | +2.5% |

As seen in Table S1, the total number of haploblocks increases substantially as sample size grows, particularly between 25 and 100 individuals, where the number more than doubles (+107%). However, beyond 100 individuals, the increase becomes much more gradual, with only a 3.7% change from 100 to 250 samples and 2.5% from 250 to 500 samples. This suggests that after a certain threshold, the software stabilizes in its haploblock detection capabilities.

#### **Table S2 Effect of sample size on genomic coverage,** displays the total genomic coverage (in base pairs) for various sample size of genotyped individuals from the DESIR cohort, with relative change indicating the variation in coverage as sample size increases

| **Sample Size** | **Coverage (bp)** | **Relative Change** |
| --- | --- | --- |
| 25 | 34,096,342 | - |
| 50 | 34,131,043 | +0.10% |
| 100 | 33,890,252 | -0.71% |
| 250 | 32,834,246 | -3.2% |
| 500 | 31,802,966 | -3.2% |

In contrast, Table S2 shows that genomic coverage remains relatively stable at smaller sample sizes but starts to decrease as the sample size increases beyond 100 individuals. The reduction in coverage suggests that the detection process is refining haploblocks, leading to finer resolution but also potentially filtering out larger blocks that may still be biologically meaningful. While larger sample sizes allow for the detection of more haploblocks, many of these additional haploblocks tend to be smaller in size. From a biological perspective, excessively small haploblocks may result from noise, sequencing artifacts, or minor variations in recombination events, making them less relevant for functional analysis. Thus, it is crucial to find a balance between accuracy and resolution—ensuring robust haploblock detection while minimizing false positives and excessive fragmentation. Based on our findings, a sample size of 100–250 individuals appears to offer an optimal trade-off: At this range, haploblock detection is stable. Larger haploblocks remain well-defined. The risk of excessive, biologically uninformative small haploblocks is minimized. Beyond 250 individuals, the increase in detected haploblocks is minor, but the proportion of smaller haploblocks rises, potentially introducing noise rather than improving accuracy. Therefore, for most studies, a sample size of around 100–250 individuals may be the most efficient choice for haploblock analysis.

**Table S3** Average haploblock size according to the sample size of genotyped individuals from the DESIR cohort: average number of SNPs per haploblock for different sample sizes, showing how haploblock size varies with increasing sample size.

| ***Sample Size*** | ***Mean size Haploblocks (# SNPs)*** |
| --- | --- |
| *25* | *33.54* |
| *50* | *35.48* |
| *100* | *27.16* |
| *250* | *25.49* |
| *500* | *25.20* |

***Speed***

To evaluate their performance further, we tested these modes in three small regions of chromosome 22 from 250 genotyped individuals of the DESIR cohort with varying SNP densities: Region 1 (13,032 SNPs, reduced to 9,303 SNPs after filtering for MAF > 0.01), Region 2 (13,255 SNPs, reduced to 8,824 SNPs after filtering for MAF > 0.01), and Region 3 (19,637 SNPs, reduced to 12,777 SNPs after filtering for MAF > 0.01).

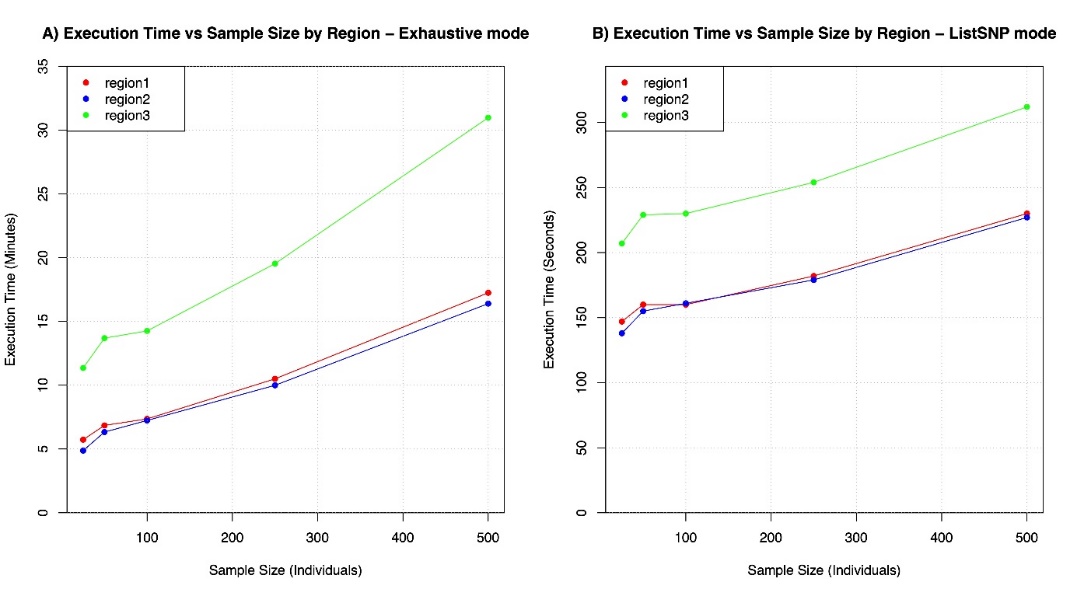

**Figure S5 -** (A) Execution Time vs Sample Size by Region in Exhaustive Mode, relationship between execution time (in minutes) and sample size (number of individuals) for three different regions (Region 1 : 9303 SNPs, Region 2: 8824 SNPs, and Region 3: 12777 SNPs) using the Exhaustive mode. (B) Execution Time vs Sample Size by Region in ListSNP Mode (SNPs = 1000), relationship between execution time (in seconds) and sample size (number of individuals) for the same three different regions (Region 1, Region 2, and Region 3) using the ListSNP mode

For the exhaustive mode, the running times were analyzed across these regions with the same sample sizes as before. As expected, the computational time increased with both the number of individuals and the SNP density of the region (Table S5; *Figure S5A*). Region 3, with the highest SNP count, consistently required the most processing time. The *ListSNP* mode’s running times were also evaluated across the same regions, with varying sample sizes and fixed SNP list sizes (50, 100, 250, 500 and 1000 SNPs). As anticipated, regions with higher SNP densities, such as Region 3, required slightly more computation time than Region 1 and Region 2. Furthermore, the computational time increased with both the sample size and the size of the predefined SNP list per region (Figure S5B, Table S5). These results are coherent with the ones already obtained for a larger region such as the chromosome 22.

**Table S4** Execution Time (in Minutes) for Different Modes with various sample sizes of genotyped individuals from the DESIR cohort (25, 50, 100, 250, and 500 individuals), comparing the Standard, ListSNP, and Exhaustive modes.

| **Individuals** | **Modes** | **Time (Minutes)** |
| --- | --- | --- |
| **25** | Standard | ~ 7 |
| **50** | Standard | ~ 11 |
| **100** | Standard | ~ 20 |
| **250** | Standard | ~ 26 |
| **500** | Standard | ~ 88 |
| **25** | ListSNP | ~ 2 |
| **50** | ListSNP | ~ 2 |
| **100** | ListSNP | ~ 2 |
| **250** | ListSNP | ~ 4 |
| **500** | ListSNP | ~ 6 |
| **25** | Exhaustive | ~ 263 |
| **50** | Exhaustive | ~ 352 |
| **100** | Exhaustive | ~ 325 |
| **250** | Exhaustive | ~ 443 |
| **500** | Exhaustive | ~ 1215 |

To further evaluate computational performance, running times were compared across different imputed chromosomes (chr1, chr6, and chr22) using the Standard mode with default parameters in the DESIR cohort. For a sample of 250 individuals, the analysis of chromosome 1 with 1,633,735 SNPs required approximately 580 minutes. When increasing the sample size to 500 individuals for chromosome 6, which contained 697,685 SNPs, runtime extended to approximately 720 minutes (~12 hours). In contrast, chromosome 22, with 125,956 SNPs, was processed in just 26 minutes for 250 individuals.

**Table S5** Running Time (in Minutes) for Exhaustive Mode for three 5 Mb Regions of Chromosome 22 with phased data (Region 1: 9303 SNPs, Region 2: 8824 SNPs, and Region 3: 12777 SNPs), according to the sample size of genotyped individuals from the DESIR cohort

| **Individuals** | **Region** | **Time (Minutes)** |
| --- | --- | --- |
| **25** | region1 | ~ 6 |
| **25** | region2 | ~ 5 |
| **25** | region3 | ~ 11 |
| **50** | region1 | ~ 7 |
| **50** | region2 | ~ 6 |
| **50** | region3 | ~ 14 |
| **100** | region1 | ~ 7 |
| **100** | region2 | ~ 7 |
| **100** | region3 | ~ 14 |
| **250** | region1 | ~ 11 |
| **250** | region2 | ~ 10 |
| **250** | region3 | ~ 20 |
| **500** | region1 | ~ 17 |
| **500** | region2 | ~ 16 |
| **500** | region3 | ~ 31 |

In this analysis, we examined the running time of the HaploExplore software in exhaustive mode across three regions of different SNP densities (Region 1: 9303 SNPs, Region 2: 8824 SNPs, and Region 3: 12777 SNPs) for varying sample sizes (25, 50, 100, 250, and 500 individuals).The running time increased with both the number of individuals and the size of the region, as expected (*Table S5*). As the number of individuals and SNPs in the region increased, so did the computational time. Region 3, with the highest SNP count, consistently required the most time for computation.

**Table S6** Average Running Time (in seconds) for ListSNP Mode across three 5 Mb Regions of chromosome 22 (Region 1: 9303 SNPs, Region 2: 8824 SNPs, and Region 3: 12777 SNPs), according to the sample size of genotyped individuals from the DESIR cohort

| **Region** | **Individuals** | **SNPs** | **Time (Seconds)** |
| --- | --- | --- | --- |
| **region1** | 25 | 50 | 7 |
| **region2** | 25 | 50 | 6 |
| **region3** | 25 | 50 | 8 |
| **region1** | 50 | 50 | 7 |
| **region2** | 50 | 50 | 7 |
| **region3** | 50 | 50 | 10 |
| **region1** | 100 | 50 | 8 |
| **region2** | 100 | 50 | 8 |
| **region3** | 100 | 50 | 11 |
| **region1** | 250 | 50 | 10 |
| **region2** | 250 | 50 | 10 |
| **region3** | 250 | 50 | 14 |
| **region1** | 500 | 50 | 18 |
| **region2** | 500 | 50 | 18 |
| **region3** | 500 | 50 | 25 |
| **region1** | 25 | 100 | 9 |
| **region2** | 25 | 100 | 9 |
| **region3** | 25 | 100 | 12 |
| **region1** | 50 | 100 | 10 |
| **region2** | 50 | 100 | 10 |
| **region3** | 50 | 100 | 14 |
| **region1** | 100 | 100 | 11 |
| **region2** | 100 | 100 | 12 |
| **region3** | 100 | 100 | 16 |
| **region1** | 250 | 100 | 15 |
| **region2** | 250 | 100 | 15 |
| **region3** | 250 | 100 | 20 |
| **region1** | 500 | 100 | 26 |
| **region2** | 500 | 100 | 25 |
| **region3** | 500 | 100 | 33 |
| **region1** | 25 | 250 | 21 |
| **region2** | 25 | 250 | 20 |
| **region3** | 25 | 250 | 30 |
| **region1** | 50 | 250 | 24 |
| **region2** | 50 | 250 | 23 |
| **region3** | 50 | 250 | 34 |
| **region1** | 100 | 250 | 25 |
| **region2** | 100 | 250 | 25 |
| **region3** | 100 | 250 | 36 |
| **region1** | 250 | 250 | 32 |
| **region2** | 250 | 250 | 32 |
| **region3** | 250 | 250 | 44 |
| **region1** | 500 | 250 | 49 |
| **region2** | 500 | 250 | 49 |
| **region3** | 500 | 250 | 65 |
| **region1** | 25 | 500 | 51 |
| **region2** | 25 | 500 | 47 |
| **region3** | 25 | 500 | 71 |
| **region1** | 50 | 500 | 56 |
| **region2** | 50 | 500 | 54 |
| **region3** | 50 | 500 | 79 |
| **region1** | 100 | 500 | 58 |
| **region2** | 100 | 500 | 58 |
| **region3** | 100 | 500 | 82 |
| **region1** | 250 | 500 | 69 |
| **region2** | 250 | 500 | 68 |
| **region3** | 250 | 500 | 96 |
| **region1** | 500 | 500 | 96 |
| **region2** | 500 | 500 | 96 |
| **region3** | 500 | 500 | 132 |
| **region1** | 25 | 1000 | 147 |
| **region2** | 25 | 1000 | 138 |
| **region3** | 25 | 1000 | 207 |
| **region1** | 50 | 1000 | 160 |
| **region2** | 50 | 1000 | 155 |
| **region3** | 50 | 1000 | 229 |
| **region1** | 100 | 1000 | 160 |
| **region2** | 100 | 1000 | 161 |
| **region3** | 100 | 1000 | 230 |
| **region1** | 250 | 1000 | 182 |
| **region2** | 250 | 1000 | 179 |
| **region3** | 250 | 1000 | 254 |
| **region1** | 500 | 1000 | 230 |
| **region2** | 500 | 1000 | 227 |
| **region3** | 500 | 1000 | 312 |

The running times for the ListSNP method were measured across three regions (Region 1, Region 2, and Region 3), with varying sample sizes and SNP numbers (Table S6).

As expected, the regions with higher SNP counts, such as Region 3 (12777 SNPs), consistently required more time than Region 1 (9303 SNPs) and Region 2 (8824 SNPs). However, since the number of SNPs was fixed at 50, 100, 250, 500 and 1000 we observe that the running time increases with both the sample size and the SNP list per region.

**Comparison with other haploblock detection softwares**

| Tool | Number of Haploblocks | Mean Haploblock Size (bp) | Max Haploblock Size (bp) | Standard Deviation (bp) | Runtime |
| --- | --- | --- | --- | --- | --- |
| HaploExplore | 547 | 56,865.19 | 574,879 | 67,606.29 | *~50 sec* |
| PLINK | 530 | 5,348.24 | 284,694 | 15,057.21 | *~2 min* |
| Big-LD | 176 | 18,032.3 | 296,637 | 38,367.54 | *~50 sec* |
| HaploBlocker | 143 | 78,703.35 | 760,929 | 94,408.67 | *~5 sec* |

**Table S7 - Haploblock Detection Results for Small Region (Region1 - 13,032 Variants of the chromosome 22 with 250 genotyped individuals from the DESIR cohort),** comparison of haploblock detection tools based on the number of identified haploblocks, mean and maximum haploblock size, standard deviation, and runtime.

In table S7, we compared the 4 software, namely HaploExplore, PLINK, Big-LD, and HaploBlocker, on a small region of chromosome 22 (13,032 variants) with 250 random individuals from the DESIR cohort, with their default settings. The results reveal notable differences, likely due to each tool’s distinct purpose and haploblock identification method. HaploBlocker, for instance, is designed to represent genetic variation in a compact set of haplotype blocks, leading to fewer but larger blocks. This is reflected in its low haploblock count (143) and high mean haploblock size (~78.7 kb), making it the most broad-scale approach. Big-LD, which detects subgroups of SNPs based on |r| correlation thresholds and partition SNPs using an interval graph, also identified relatively few blocks (176), with intermediate block sizes (~18 kb mean). In contrast, PLINK applied Gabriel’s method using D’ confidence intervals, leading to smaller and more numerous haploblocks (530 blocks, ~5.3 kb mean size). HaploExplore, designed for MiA-haploblock detection, produced 547 haploblocks, with a mean block size (~56.9 kb) notably larger than PLINK’s but smaller than the one of HaploBlocker. Its approach, which orders SNPs by MAF and incorporates LD thresholds and carrier percentage, allows for a flexible and biologically meaningful haploblock definition, particularly suited for minor allele-driven associations. Computational efficiency also varied: PLINK and HaploExplore completed in ~2 minutes, Big-LD in ~50 seconds, and HaploBlocker in just ~5 seconds, highlighting different algorithmic complexities, due to the different haploblock construction (Table 1).

**Table S8** Comparison with other software: running time obtained for the haploblock analysis of 250 genotyped individuals from the DESIR cohort in full chromosome 22

| **Tool** | **Runtime (chr22)** |
| --- | --- |
| HaploExplore | ~26 min |
| Plink | ~36 min |
| Big-LD | ~7 min |
| HaploBlocker | ~2 min |
